# Single Molecule Visualization of Cardiac Myosin-Binding Protein C N-terminal Fragments Interacting with Thin Filaments: Mechanisms of Calcium Sensitization

**DOI:** 10.1101/421776

**Authors:** Alessio V. Inchingolo, Samantha Beck Previs, Michael J. Previs, David M. Warshaw, Neil M. Kad

**Affiliations:** School of Biological Sciences, University of Kent, Canterbury, CT2 7NH, UK.; Department of Molecular Physiology and Biophysics, University of Vermont, Burlington, Vermont 05405 USA

**Keywords:** Actin, cMyBP-C, muscle, contractility, cardiomyopathy, diffusive motion

## Abstract

Cardiac muscle contraction is activated by calcium binding to troponin and the consequent motion of tropomyosin on actin within the sarcomere. These movements permit myosin binding, filament sliding and motion generation. One potential mechanism by which the N-terminal domains of cardiac myosin-binding protein C (cMyBP-C) play a modulatory role in this activation process is by cMyBP-C binding directly to the actin-thin filament at low calcium levels to enhance the movement of tropomyosin. To determine the molecular mechanisms by which cMyBP-C enhances myosin recruitment to the actin-thin filament, we directly visualized fluorescently-labelled cMyBP-C N-terminal fragments and GFP-labelled myosin molecules binding to suspended actin-thin filaments in a fluorescence-based single molecule microscopy assay. Binding of the C0C3 N-terminal cMyBP-C fragment to the thin filament enhanced myosin association at low calcium levels. However, at high calcium levels, C0C3 bound cooperatively, blocking myosin binding. Dynamic imaging of thin filament-bound Cy3-C0C3 molecules demonstrated that these fragments diffuse along the thin filament before statically binding, suggesting a mechanism that utilizes a weak-binding mode to search for access to the thin filament and a tight-binding mode to sensitize the thin filament to calcium and thus, enhance myosin binding. Although shorter N-terminal fragments (Cy3-C0C1 and Cy3-C0C1f) bound to the thin filaments and displayed modes of motion on the thin filament similar to that of the Cy3-C0C3 fragment, the shorter fragments were unable to sensitize the thin filament. Therefore, the longer N-terminal fragment (C0C3) must possess the requisite domains needed to bind specifically to the thin filament in order for the cMyBP-C N terminus to modulate cardiac contractility.

## Introduction

Cardiac myosin-binding protein-C (cMyBP-C) modulates cardiac contraction at the level of the sarcomere. Mutations in the cMyBP-C gene are a major cause of hypertrophic cardiomyopathy (1), a disease that affects up to 1 in 200 people (2) and is the leading cause of sudden cardiac death in young adults (1, 3). cMyBP-C is composed of 11 subdomains from C0 to C10 (Fig. 1a), 8 Ig-like and 3 Fn(III)-like with a proline-alanine rich region between C0 and C1, and a linker between C1 and C2 (i.e. M-domain) (4). Each cMyBP-C molecule is tethered through its C-terminus (5) to the myosin thick filament (6), while its N-terminal domains are free to contact either the myosin head region or the actin thin filament (7) (Fig. 1a). Specifically, cMyBP-C interactions with myosin have been shown to occur through the S2 region (8-11), the myosin regulatory light chain (RLC) (12) and the S1 head (13), which may serve to stabilize myosin’s super relaxed state (14-16) through interactions with the recently defined myosin mesa region (13, 17). In addition to myosin-binding, cMyBP-C interacts with actin via its N-terminal domains (C0-C2) (18-21). These binding partner interactions are believed to functionally impact cardiac contractility. For example, cMyBP-C knockout mouse cardiomyocytes (22) demonstrate a decrease in calcium sensitivity while N-terminal fragments infused into skinned fibres (11) result in an increase in calcium sensitivity. Accordingly, fluorescence assays in muscle cells (10) have demonstrated that cMyBP-C N-terminal fragments activate muscle at low calcium levels and inhibit maximal activity at high calcium, suggesting dual roles in modulating contraction *in vivo*.

**Figure 1.**
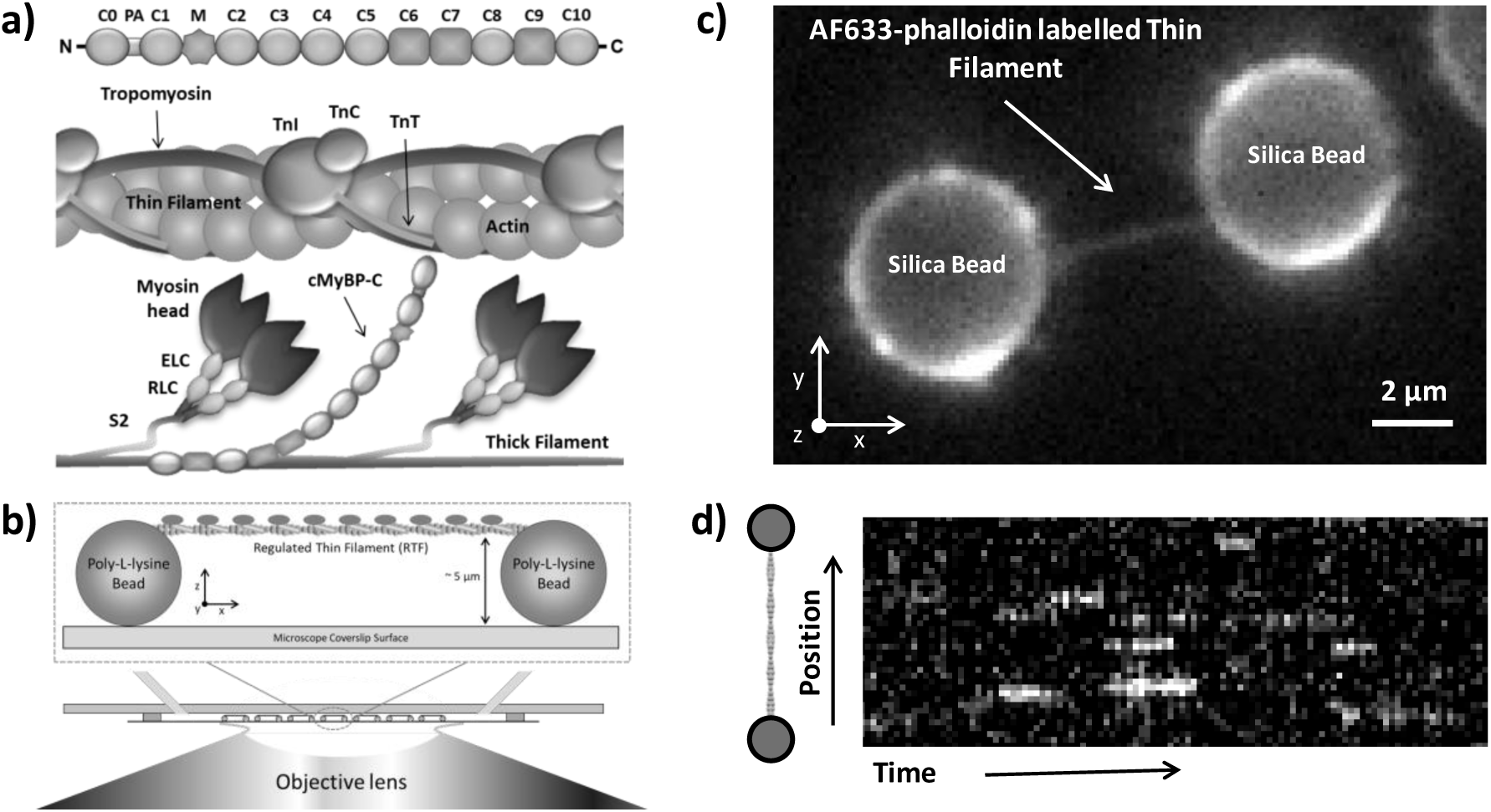
Thin filaments in vivo and in the single molecule tightrope assay: a) The subdomain structure of cMyBP-C and its spatial relationship to the thin filament (actin decorated with tropomyosin and the troponin complex TnT, TnI and TnC) and myosin-containing thick filament in the sarcomere. b) The thin filament tightrope assay. Regulated thin filaments are suspended between surface adhered beads using a microfluidic device and visualised using a high NA objective lens. c) A regulated thin filament labelled with AF633-phalloidin can be visualised suspended between two beads. d) Representative kymograph of interactions between 15 nM S1-GFP and a thin filament tightrope at high calcium (pCa 4) and 0.1 μM ATP. Each x-axis pixel corresponds to a frame (300 ms), and each pixel on the y axis corresponds to 126.4 nm along the tightrope.

Mechanistically, muscle activation occurs by calcium-dependent shifts in the position of tropomyosin on the thin filament (23, 24). Tropomyosin binds axially along the thin filament (Fig. 1a) and shifts azimuthally when calcium binds to the tropomyosin-associated troponin protein complex. Three positions of tropomyosin have been structurally (24) and functionally (25) defined; blocked (sterically preventing myosin binding), closed (myosin can bind weakly) and open (myosin binding is unperturbed). It has been proposed that the cMyBP-C N terminus could sensitize the thin filament to calcium through the displacement of tropomyosin from its blocked to its closed position, to enhance myosin binding (21). The function of the cMyBP-C N-terminal sub-domains C0C3, C0C1 and C0C1f (i.e., C0C1 including the first 17 M-domain residues) have been extensively characterized for their ability to sensitize the thin filament to calcium using *in vitro* motility (18, 26, 27), ATPase assays (19), and electron micrographic studies (18, 21, 28). Our previous studies and that of others show that all three bind to actin, but only the C0C3 fragment can both activate thin filament sliding at low (0.1 μM) calcium and inhibit sliding at high (100 μM) calcium levels. In contrast, the shortest fragment, the murine C0C1, does neither, while the C0C1f with its additional 17-residues of the M-domain can only inhibit thin filament sliding at high calcium (18). These differential functions may allow assignment of specific functional capacities to cMyBP-C’s N-terminal domains.

To define the molecular basis of thin filament activation by cMyBP-C, we used a single molecule suspended thin filament assay (Fig. 1b), in which we could directly observe the calcium-dependent binding of individual fluorescently labelled myosin-S1 molecules to the thin filament (29). Using all three murine cMyBP-C N-terminal fragments (Fig. 2b): C0C3, C0C1 and C0C1f, we show that cMyBP-C’s N terminus (C0C3) can maximally activate the thin filament, permitting myosin-S1 to bind even at low calcium. However, at high calcium with the thin filament maximally activated, cMyBP-C can decrease myosin-S1 binding. By imaging fluorescently labelled cMyBP-C N-terminal fragments, we observed these molecules individually binding directly to the thin filament. Once bound, the cMyBP-C fragments both diffuse along the thin filament and bind statically, with enhanced static binding observed only for the C0C3 fragment at high calcium levels that maximally activate the thin filament. These fragment binding and motion behaviours on the thin filament provide direct mechanistic insight into how cMyBP-C achieves both calcium sensitization at low calcium and reduced activation at high calcium, and thus modulation of cardiac contractility.

**Figure 2.**
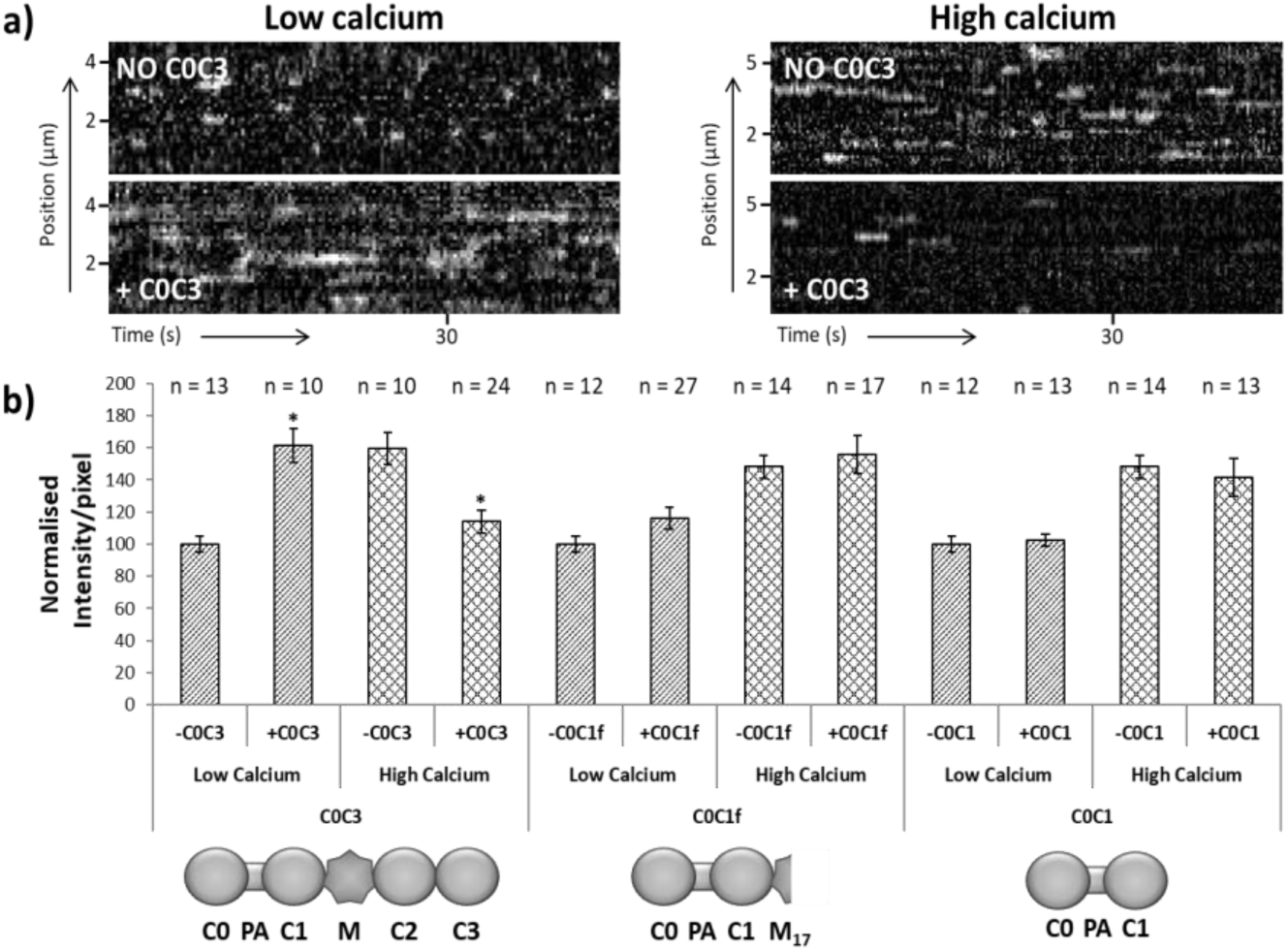
The calcium-dependent effects of cMyBP-C fragments on S1-GFP binding to regulated thin filaments: a) Representative kymographs of S1-GFP binding to thin filaments in the absence and presence of C0C3, at low and high calcium, collected at 3.3 frames per second. b) Bar chart of the average intensity/pixel ratio in the kymographs of S1-GFP binding in the absence and presence of different cMyBP-C N-terminal fragments; the values have been normalised to each respective control experiment (i.e. no fragment present at low [Ca^2+^]). Data were collected using 15 nM S1-GFP w/or w/o 1 μM of fragment, with n values representing the number of tightropes imaged per condition. Results showed a significant increase (61.2% ± 11.7 (SEM), p < 0.001) in S1-GFP binding in the presence of C0C3 at low calcium and a significant reduction (45.5% ± 11.9 (SEM), p = 0.009) at high calcium; no significant difference is seen when using the shorter fragment C0C1f (16.0% ± 8.2 (SEM), p = 0.1390 at low calcium and 7.5% ± 13.8 (SEM), p = 0.6126 at high calcium) and C0C1 at low (2.4% ± 6.2 (SEM), p = 0.7072) and high calcium (−6.8% ± 13.7(SEM), p = 0.6189).

## Methods

### Protein purification

Actin and myosin were purified from chicken pectoralis (30), while recombinant human cardiac tropomyosin and troponin were expressed in *E. coli* and further purified as described previously (31). Murine cMyBP-C fragments were bacterially-expressed with a C-terminal His-tag for purification using a nickel affinity column. Myosin-S1 was produced from whole myosin by papain digestion (32), and labelled with eGFP by exchange of the endogenous regulatory light chain (RLC) with recombinantly expressed eGFP-tagged RLC (29); this construct is termed S1-GFP. Using the proteolytically derived single-headed S1 construct, instead of full-length myosin or heavy meromyosin, simplified interpretation and quantification of the data. Thin filaments were fully reconstituted by incubating actin, tropomyosin (Tm) and troponin (Tn) overnight at 4°C in 4 mM Imidazole pH 7, 2 mM MgCl_2_, 1 mM DTT, 0.5 mM ATP (33) with a molar ratio actin:Tm:Tn of 2:0.5:0.25. In all experiments, thin filaments were labelled with AF633-phalloidin (Invitrogen, A22284) for imaging in the flow chamber. For experiments involving cMyBP-C N-terminal fragments binding to thin filaments, the N-terminal fragments (i.e., C0C3, C0C1f and C0C1, see Fig. 2b) were fluorescently labelled with a maleimide-Cy3 (GE Healthcare, Amersham, PA23031) as in (34), achieving ^~^30% labelling efficiency. Based on the C0C3 structure, there is only one surface exposed reactive cysteine (C248) and thus only one Cy3 dye per labelled N-terminal fragment (34).

### The thin filament tightrope assay

To nullify any potential surface interference and to permit full, three-dimensional accessibility to all myosin and cMyBP-C binding sites on the thin filament, we suspended thin filaments between surfaced adhered beads to create tightropes (Fig. 1b) as described in detail previously (29). Briefly, silica beads were functionalised with 340 μg/ml poly-L-lysine and adhered to a glass coverslip by infusion into a microfluidic flow cell. To generate tightropes, 500 nM thin filaments were passed across the silica beads multiple times back and forth using a syringe pump. Imaging was performed using a custom-built oblique angle fluorescence (OAF) microscope (35) that excites the sample with a continuous wave 20 mW 488 nm DPSS laser (JDSU) focused off-centre at the back-focal plane of a 100x objective (1.45NA) to achieve obliquely angled illumination. We detected images (Fig. 1c) through an Optosplit III (Cairn Research, UK) projected onto a Hamamatsu OrcaFlash 4.2 camera.

### Data collection and analysis

All experiments were performed in either high (100 μM, pCa 4) or low (0.1 μM, pCa 7) calcium buffers of 25 mM imidazole pH 7.4, 4 mM MgCl_2_, 1 mM EGTA, and KCl adjusted to a final ionic strength of 51 mM using MaxChelator (36). Ten or more tightropes were imaged per condition (details of the conditions are provided in the Results). Videos of S1-GFP binding to thin filaments were acquired at 3 frames per second (fps) for up to 1 minute, while videos of Cy3-labelled cMyBP-C fragments were acquired at 1 fps for up to 5 minutes. To determine the spatial and temporal dynamics of S1-GFP or Cy3-labelled cMyBP-C fragments interacting with thin filaments, we transformed movies into kymographs where fluorophore position was plotted along the y-axis and time on the x-axis (Fig. 1d). Analysis of bound fluorophore position was performed using custom written macros in ImageJ available online (http://kadlab.mechanicsanddynamics.com). To measure the extent of S1-GFP binding we integrated the fluorescence intensity over the entire kymograph and normalised to the total number of pixels. All statistical p-values mentioned were calculated using an unpaired t test, while all the SEM were calculated according to 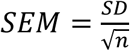 and propagated when necessary.

## Results

### The effect of cMyBP-C N-terminal domains on S1-GFP myosin binding to the thin filament

We studied the ability of each of the three N-terminal fragments C0C3, C0C1f and C0C1 (Fig. 2) to sensitize the thin filament to calcium (18-21), as evidenced by S1-GFP’s binding to thin filaments. Using 1 μM unlabelled cMyBP-C N-terminal fragments, we determined the level of S1-GFP (15 nM) binding to the thin filament at low and high calcium (0.1 μM and 100 μM, respectively). The fragment concentration was chosen based on past motility studies in which 1 μM C0C3 both sensitized the thin filament to calcium and inhibited thin filament motility at maximally activating calcium concentrations (37). Figure 2a shows representative kymographs for S1-GFP binding. At low calcium in the absence of C0C3, the kymograph (Fig. 2a, upper left) shows very few S1-GFP binding events, consistent with the thin filament not being fully activated, as previously described (29). At high calcium in the absence of C0C3, the thin filament is activated as evidenced by significant S1-GFP binding to the thin filament (Fig. 2a, upper right). In the presence of C0C3 the situation is reversed. At low calcium, considerable S1-GFP binding is observed (Fig. 2a lower left), whereas at high calcium these interactions are rare (Fig. 2a, lower right). To quantify changes in S1-GFP binding, we calculated the average fluorescence intensity per pixel for each kymograph, averaged over at least 3 kymographs per tightrope. Minimally, 10 tightropes were observed under the same experimental condition (Fig. 2b: n, total tightrope number). To compare the effects of a given cMyBP-C fragment on S1-GFP binding, the average fluorescence intensity per pixel was normalised to that at low calcium concentration in the absence of fragments (Fig. 2b). At low calcium, addition of C0C3 significantly increased (p < 0.0001) S1-GFP binding (61.2% ± 11.7 (SEM)) versus no fragment; equivalent to that observed at high calcium in the absence of C0C3 (59.5% ± 10.9 (SEM)). However, addition of C0C3 at high calcium, significantly (p = 0.009) reduced S1-GFP binding by 45.5% ± 11.9 (SEM) versus the absence of C0C3 at high calcium. Interestingly, no significant difference in S1-GFP binding was seen as a result of the introduction of either shorter fragment, C0C1f or C0C1, at both low and high calcium. These results suggest that the shorter fragments are missing critical domains essential for modulating S1-GFP binding to the thin filament in a calcium-dependent manner.

### Imaging N-terminal cMyBP-C fragments binding to the thin filament

As described above, only the C0C3 fragment affected S1-GFP binding to the thin filament in a calcium-dependent manner. Could the differential effects seen for the C0C3 fragment compared to the shorter C0C1f and C0C1 fragments be related to a difference in their binding to the thin filament? Therefore, we imaged 20 nM Cy3-tagged C0C3, C0C1f and C0C1 interacting with thin filaments at both low and high calcium. Figure 3 shows clear binding of all three fragments to the thin filaments using long exposure (1 sec) still images. These images were analysed to determine the average number of fragment molecules bound per micron of thin filament. To ensure a more accurate determination of closely spaced fluorescent fragments that could not be spatially resolved, we used fluorescent spot intensity to estimate the number of molecules per spot. Firstly, we determined the fluorescence intensity for single Cy3-C0C3 molecules from thin filament binding images at low calcium (Fig. 3a, left panel), where fluorescent spots were easily discernible. Even in these conditions there was considerable variance in spot intensity, suggesting that such spots were due to closely spaced, neighbouring Cy3-C0C3 molecules. To extract the fluorescent intensity of a single molecule, we fitted the distribution of intensities at low calcium to multiple Gaussians (29, 38) as detailed in the supplementary information and Fig. S1. Using the fluorescence intensity of a single Cy3, we converted each fluorescence spot intensity along a tightrope into the number of bound molecules and used this to calculate the average number of cMyBP-C fragments bound per micron of thin filament. For Cy3-C0C3 (Fig. 3a & d), there was a >5-fold increase (p < 0.0001) in binding at high versus low calcium (5.69 ± 0.55 molecules/μm (SEM, n = 28 tightropes) vs. 1.0 ± 0.12 molecules/μm (SEM, n = 50 tightropes), respectively). This was not the case for Cy3-C0C1f (Fig. 3b & d), where no significant (p = 0.0523) change in decoration of thin filaments (in molecules/μm) was observed between low and high calcium (1.14 ± 0.11 (SEM, n = 94 tightropes) vs. 1.61 ± 0.21 (SEM, n = 99 tightropes) molecules/μm, respectively). However, more Cy3-C0C1 (Fig. 3c & d) bound to thin filaments (p = 0.0055) at low (2.76 ± 0.37 (SEM, n = 68 tightropes) molecules/μm) versus high calcium (0.73 ± 0.23 (SEM, n = 19 tightropes) molecules/μm). A small (p < 0.0001) but significant increase in Cy3-C0C1 decoration of the thin filaments was observed at low calcium compared with both Cy3-C0C3 and Cy3-C0C1f. In summary, neither C0C1 or C0C1f (at high or low calcium) bound substantially differently to C0C3 at low calcium. Only C0C3 bound strongly to the thin filaments at high calcium.

**Figure 3.**
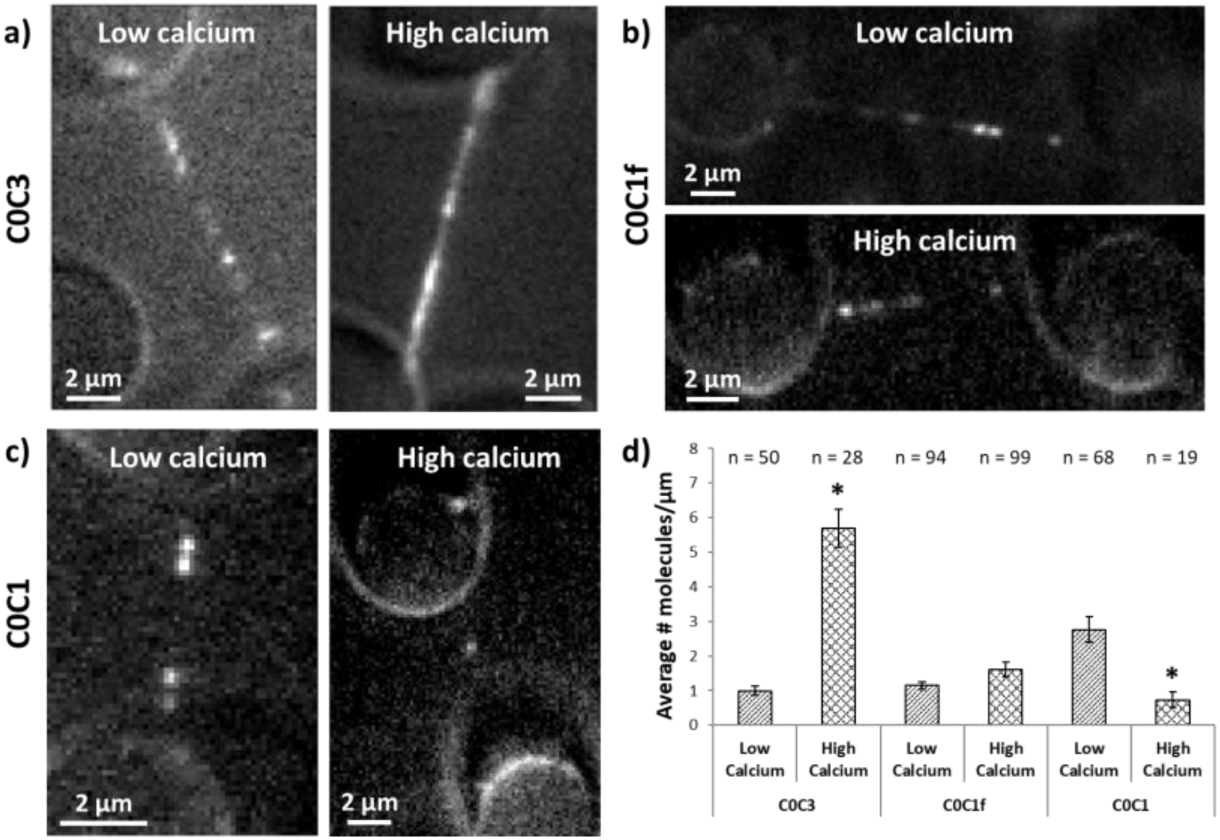
Fluorescent cMyBP-C N-terminal fragments associate with thin filament tightropes: Representative images of tightropes decorated using 20 nM Cy3-tagged a) C0C3, b) C0C1f and c) C0C1, in low and high calcium conditions. d) Bar chart indicating the average number ± SEM of Cy3-tagged N-terminal cMyBP-C fragments bound per μm of tightrope. n indicates number of tightropes imaged with significance of p<0.01 identified by an ^∗^.

The binding of Cy3-C0C3 at high calcium to the thin filament appeared as clusters suggesting the existence of cooperativity. To quantify this, we determined the number of molecules per fluorescent spot for Cy3-C0C3; the only fragment that showed increased binding at high calcium. Figure 4 shows the distribution of number of molecules within each fluorescent spot or cluster on a thin filament for Cy3-C0C3 at low and high calcium. At low calcium, the majority of fluorescent spots contain only a single Cy3-C0C3; however, at high calcium the distribution is quite broad with its peak at four Cy3-C0C3 molecules per spot. These plots can be described by polymer association probabilities that are modified by the availability of actin. We used the Flory (also termed Flory-Schulz) distribution that describes the weight fraction of a polymer through the probability of monomer association (39), as described by the following equation:

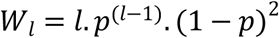

where *W_l_* is the weight fraction and *l* is the number of molecules in the cluster. *p* is the extent of reaction and is governed by the affinity of Cy3-C0C3 for actin:

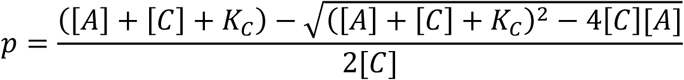

**Figure 4.**
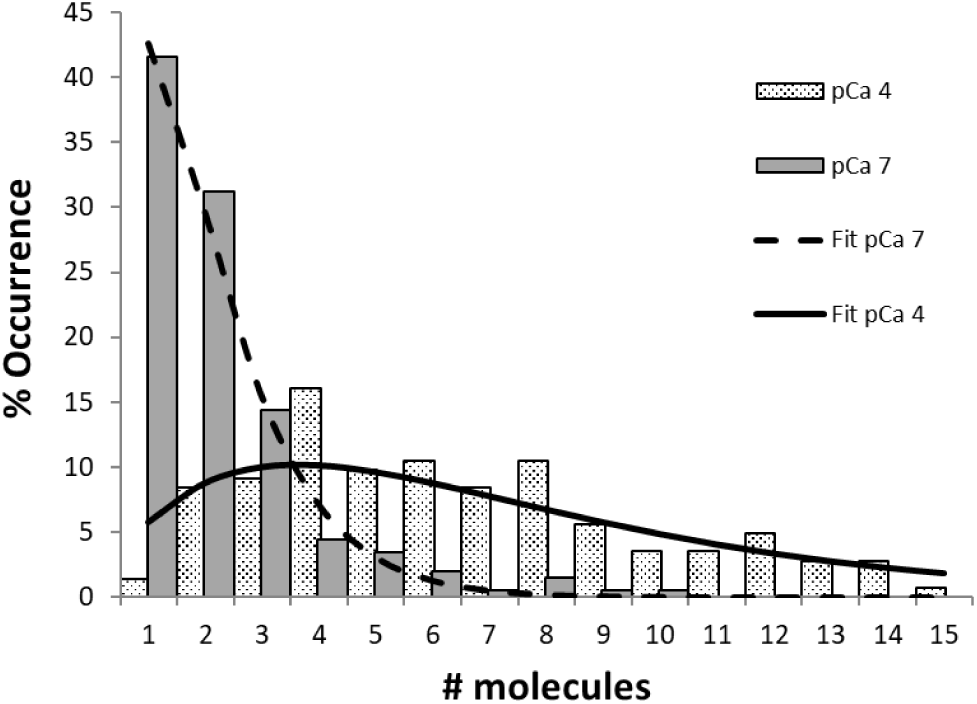
Histogram of cMyBP-C cluster sizes: The number of fluorescent molecules bound to the thin filament tightropes were determined based on the fluorescence intensity of a single Cy3-labelled C0C3 (see Fig. S1). Plotted at low calcium (solid bars) and high calcium (dotted bars), data are fitted simultaneously to a Flory distribution (see main text) yielding excellent fits with fit parameters provided in the main text.

Where *A* is the actin concentration, *C* is the Cy3-C0C3 concentration and *K_c_* is the affinity of the fragment for actin. However, since the availability of actin changes with calcium concentration we nested the following term for [A]:

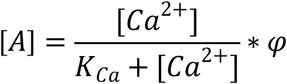

where *K_Ca_* is the thin filament calcium affinity and *Ca^2+^* is the calcium concentration. *φ* is the apparent concentration of actin in the flowcell.

The overlaid fit in Figure 4 was performed in Microsoft Excel using the GRG Nonlinear multistart fitting engine to globally fit both the high and low calcium conditions simultaneously. The affinity (*K_ca_*) of calcium binding to thin filaments was determined to be 0.46 μM (pCa_50_ of ^~^6.34), in full agreement with previous studies (40), *φ* was fitted as 400 nM consistent with the 500 nM used to create the tightropes, while the Cy3-C0C3 affinity for actin *K_c_*, was found to be 0.12 μM. The increase in number of molecules per cluster is clearly seen at high calcium, indicating cooperative binding. This is confirmed by the fit to the Flory polymer distribution model.

### N-terminal cMyBP-C fragments display two modes of motion on thin filaments

Unexpectedly, in addition to N-terminal fragments binding statically to thin filaments (Fig. 5a, horizontal trace (arrowhead)), a proportion of the fragments appeared to diffuse randomly along the thin filament (Fig. 5a, trace with y-axis movement (star)). This behaviour was calcium dependent for Cy3-C0C3 molecules; the proportion diffusing decreased significantly (p = 0.029) from 19.4% ± 4.6 (SEM) at low calcium to 6.0% ± 1.8 (SEM) at high calcium (Fig. 5b). By comparison, Cy3-C0C1f showed no significant difference (p = 0.2025) in the proportion of diffusive molecules at both low and high calcium, 27.3% ± 2.2 (SEM) and 32.3% ± 3.1 (SEM), respectively. This was also the case for Cy3-C0C1 (24.5% ± 6.4 (SEM) vs. 47.7% ± 26.2 (SEM) at low and high calcium, respectively; p = 0.2542).

**Figure 5.**
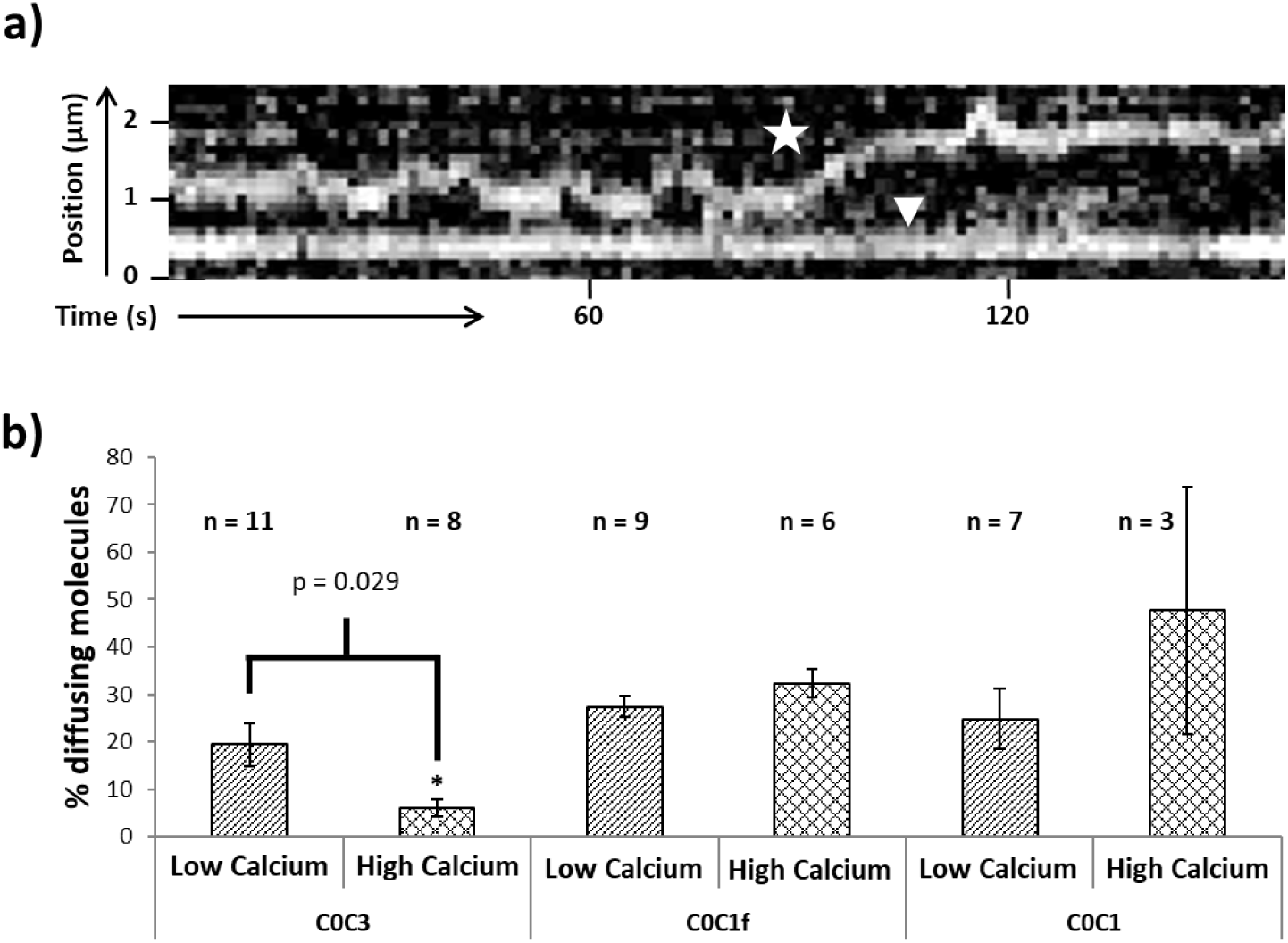
cMyBP-C fragments diffuse on and statically bind to thin filament tightropes: a) An example of a kymograph representing the two types of interaction seen with cMyBP-C (using Cy3-C0C3 as an example). Diffusion on the thin filament tightrope is indicated by star and a stationary molecule is indicated with an arrowhead. b) A bar chart providing the mean fraction ± SEM of diffusing molecules measured at low and high calcium concentrations. n values refer to the number of experiments with significance of p<0.05 identified by an ^∗^. Data were collected using 20 nM of either C0C3, C0C1f and C0C1 at low calcium; at high calcium, to compensate for decoration levels that enable individual molecules to be distinguished, 5 nM C0C3 and C0C1f or 20 nM C0C1 was used.

We calculated the diffusional characteristics summarised in Table 1 using mean squared displacement (*MSD*):

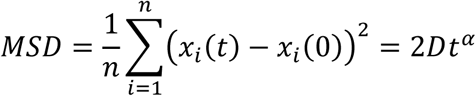

where *D* is the diffusion constant of the molecule, *t* is the analysis time window, *x_i_*(*t*) is the position at time t, *x_i_*(0) is the starting reference position and *α* is a coefficient that denotes the nature of the diffusion, where values close to 1 indicate random diffusion (41). Table 1 summarises the values we obtained for *D* and *α* from this analysis.

**Table 1:** Diffusional characteristics for N-terminal cMyBP-C constructs versus calcium concentration.

|  | Cy3-COC3 |  |  |  | Cy3-COC1f |  |  |  | Cy3-COC1* |  |
| --- | --- | --- | --- | --- | --- | --- | --- | --- | --- | --- |
| | 0.1 $\mu\text{M}$ $\text{Ca}^{2+}$ | | 100 $\mu\text{M}$ $\text{Ca}^{2+}$ | | 0.1 $\mu\text{M}$ $\text{Ca}^{2+}$ | | 100 $\mu\text{M}$ $\text{Ca}^{2+}$ | | 0.1 $\mu\text{M}$ $\text{Ca}^{2+}$ | |
| | D ($\mu\text{m}^2/\text{s}$)<br>$\times 10^{-3}$ | $\alpha$ | D ($\mu\text{m}^2/\text{s}$)<br>$\times 10^{-3}$ | $\alpha$ | D ($\mu\text{m}^2/\text{s}$)<br>$\times 10^{-3}$ | $\alpha$ | D ($\mu\text{m}^2/\text{s}$)<br>$\times 10^{-3}$ | $\alpha$ | D ($\mu\text{m}^2/\text{s}$)<br>$\times 10^{-3}$ | $\alpha$ |
| Mean | 6.28 | 0.98 | 0.68 | 1.36 | 4.06 | 1.00 | 2.95 | 0.88 | 1.13 | 0.71 |
| SEM | 1.24 | 0.07 | 0.21 | 0.16 | 0.90 | 0.07 | 1.152 | 0.08 | 0.40 | 0.09 |
| n | 32 | 33 | 5 | 6 | 39 | 39 | 15 | 15 | 9 | 12 |

\*Due to insufficient binding, diffusion data were not acquired for COC1 at 100 $\mu\text{M}$ $\text{Ca}^{2+}$

Diffusion along a lattice can be described as multiple discrete steps with an average stepping rate. This definition allows us to calculate the average step size of the fragments on the thin filament, provided the stepping rate is known. We can calculate the stepping rate from attached lifetimes obtained from laser trap studies (42) that indicated C0C3, C0C1f and C0C1 attach with two lifetime populations, <30 ms and >200 ms. Given that the longer events were ^~^60 times less frequently observed, we used the short lifetime to calculate the step size for each fragment at low calcium, according to equations below (35):

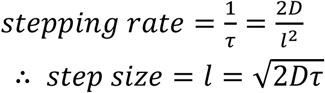

where *τ* is the attached lifetime, *D* is the diffusion constant and *l* is the average step size. This provides an average step size of 17 ± 3 nm (SD) for Cy3-C0C3, 12 ± 2 nm (SD) for Cy3-C0C1f and 7 ± 1 nm (SD) for Cy3-C0C1. Due to the method of calculating these values it was not possible to generate a significance test, therefore these values provide a range of step size from ^~^1 to 3 actin monomers.

## Discussion

cMyBP-C plays a critical role in modulating cardiac contractility (43, 44), as evidenced by high prevalence of cMyBP-C mutations in hypertrophic cardiomyopathy (45). One way for cMyBP-C to modulate cardiac contractility is to alter the sensitivity of the thin filament to calcium (10, 18, 27, 43), as reported here. To determine the underlying mechanism by which cMyBP-C enhances thin filament activation at low calcium levels (18, 26), we directly visualised the impact of cMyBP-C N-terminal fragments on the binding of individual myosin (S1-GFP) molecules along a single suspended actin-thin filament. Specifically, we observed that the C0C3 N-terminal fragment enhanced myosin binding to thin filaments at low calcium (i.e. increased calcium sensitivity) and reduced myosin binding at high calcium. These results are consistent with our previous *in vitro* motility studies and that of others (18, 26), showing that cMyBP-C’s N terminus is sufficient to sensitize the thin filament to calcium and to modulate thin filament sliding velocities at high calcium. The binding of Cy3-C0C3 itself to the thin filament is seen to be cooperative and that this binding most likely displaces tropomyosin to a position that facilitates myosin binding (10, 18, 20). Surprisingly, ^~^20% of total Cy3-C0C3 binding events show diffusive behaviour on the thin filament at low calcium, which may reflect the behaviour of cMyBP-C’s N terminus within the sarcomere as it searches for its binding site on the thin filament.

### Specific cMyBP-C N-terminal domains are necessary for thin filament activation

Activation of the thin filament at high calcium triggered significant S1-GFP binding, as expected (Fig. 2). Interestingly, the C0C3 N-terminal fragment sensitized the thin filament to calcium so that at low calcium, S1-GFP binding was equivalent to that for a fully calcium-activated thin filament (Fig. 2). The shorter N-terminal fragments (C0C1f and C0C1), with the C2 and C3 domains and a portion or the entire M-domain removed, were still capable of binding to thin filaments (Fig. 3). However, these fragments failed to sensitize the thin filament at low calcium (Fig. 2), measured as no change in S1-GFP binding in their presence. The failure of these shorter, murine cMyBP-C N-terminal fragments to sensitize the thin filament agrees with ATPase (19), *in vitro* motility (18, 26), and fragment-infused fibre data (11), suggesting that murine N-terminal fragments require at least C0-C2 to observe maximal thin filament calcium sensitization as compared to the human N-terminal cMyBP-C fragments, where C0C1 and C1 alone are sufficient to sensitize the thin filament to calcium (19, 20, 46). This species difference has been attributed to alternate sequences in the Pro-Ala linker between C0 and C1 as well as C1 itself (47).

The three-state model for thin filament regulation (25) provides a useful mechanistic context for how C0C3 sensitizes the thin filament to calcium. Specifically, calcium binding to the thin filament permits tropomyosin movement on actin to expose myosin binding sites, shifting tropomyosin from the “blocked” to the “closed” position. As a result of myosin binding, tropomyosin is further displaced to the “open” or “myosin” position (24), therefore propagating the binding of additional myosins (29). By C0C3 binding to the thin filament at low calcium, this N-terminal fragment may similarly stabilize tropomyosin in the closed position as does calcium alone, allowing myosin binding to bind, as supported by structural studies of tropomyosin movement caused by C0C2 binding to thin filaments (18).

### Modes of cMyBP-C N-terminal interactions with the thin filament

The modular architecture of cMyBP-C provides the potential to structurally compartmentalize its functional capacities. Even though the very N-terminal domains, C0C1 and C0C1f, bind to actin (20, 21, 48, 49) as we have also shown previously (50), only C0C3 possesses the ability to sensitize the thin filament to low calcium while decreasing activation levels at high calcium. Unique to our study is the observation that upon binding of these N-terminal fragments to the thin filament, the fragments either remain stationary or undergo a diffusive-like movement along the thin filament. Specifically, for C0C3, how do these two distinct modes of interaction (i.e., 20% diffusing and 80% stationary) contribute to the thin filament sensitization at low calcium? Multiple sites of N-terminal fragment interactions with the thin filament have been proposed based on actin cross-linking (51), fragment binding to actin in the laser trap (42), and electron microscopy studies (18, 20, 21, 49). At low calcium, tropomyosin on the thin filament should be in a dynamic equilibrium between the blocked and closed states (52). Presumably, the stationary Cy3-C0C3 molecules are stereo-specifically bound to the thin filament through multiple binding sites, which could effectively shift tropomyosin’s equilibrium position to being predominantly closed and thus sensitize the thin filament to calcium. Although at low calcium, the Cy3-C0C1 and Cy3-C0C1f have the same distribution of diffusing and stationary molecules, therefore the stationary fraction are not bound in the same manner as Cy3-C0C3, given the inability of these shorter fragments to sensitize the thin filament for myosin binding. As individual Ig-domains, both C0 and C1 can bind to the thin filament in several different configurations (20), which is not the case for longer C0C2 and C0C3 fragments (21, 53). These structural studies support our proposal that the stationary Cy3-C0C3 molecules are the result of multiple sites of interaction with the thin filament, necessary to achieve calcium sensitization.

For the 20% of the Cy3-C0C3 fragments that diffuse at low calcium, these fragments must weakly interact with the thin filament with a diffusional step size that we estimate to be between ^~^7 to ^~^17 nm, consistent with 1-3 actin monomers. This diffusive mode of motion may allow the N terminus of cMyBP-C to maintain contact with and scan the thin filament until such time that specific cMyBP-C binding sites are exposed on actin as tropomyosin undergoes its dynamic equilibrium between blocked and closed positions on the thin filament. Once statically bound to these sites, cMyBP-C may keep tropomyosin in the closed position (18, 25), thus, sensitizing the thin filament to calcium. This may also explain why at high calcium where the thin filament is predominantly in the closed state (25), Cy3-C0C3 binding to the thin filament is significantly increased (Fig. 3d) and almost entirely stationary (Fig. 5b) as cMyBP-C binding sites on actin would be readily exposed.

The shift from a diffusive to a stationary mode may also underlie the apparent cooperative Cy3-C0C3 binding observed at high calcium, (Figs. 3, 4), which was not seen with the shorter C0C1f fragment (Fig. 4). The clustering of C0C3 may compete with myosin for actin binding sites, as proposed in structural studies (18, 20, 21, 49) and co-sedimentation competition studies (54), explaining the reduction in myosin binding observed on tightropes at high calcium (Fig. 2). Therefore, competitive inhibition of myosin binding by the N-terminus of cMyBP-C may be one contributing factor to the reduced thin filament velocity observed both in the C-zone of native thick filaments (55) and in the motility assay at high calcium (18, 26). Is *in vivo* clustering of cMyBP-C on the thin filament possible? cMyBP-C is positioned every ^~^43 nm along the thick filament within the C-zone. Since each thin filament in the muscle lattice is surrounded by three thick filaments, the individual cMyBP-C N termini from these thick filaments could cluster at a common point on the thin filament. In addition, HCM mutations of MyBP-C are known to lead to truncations (56), which could potentially compete with myosin as previously suggested (22). Fluorescence detection of Cy3-C0C3 binding and clustering at low versus high calcium enabled us to use a polymer growth model to estimate the affinity of these cMyBP-C fragments for thin filaments as 0.12 μM. This value is ^~^20-fold tighter than the ^~^2 μM reported for unregulated actin from ATPase inhibition studies (19) and up to 50-fold tighter than the affinities reported for mouse C0C2 using actin co-sedimentation assays (51, 57). The studies reporting the lower binding affinities (51) used bare actin, whereas thin filaments were used in the present study. Recent findings suggest that tropomyosin may bind cMyBP-C’s N terminus (58), therefore, tropomyosin itself could act as a cofactor in the cooperative binding of C0C3.

## Conclusion

In this paper, we addressed the nature of thin filament activation/inhibition by the N terminus of cMyBP-C. Using a newly developed single molecule imaging assay, we propose a mechanism by which cMyBP-C interacts with thin filaments using two different binding regimes in relaxing and activating calcium conditions. At low calcium (relaxing conditions), cMyBP-C senses the thin filament for changes in activation state due to positional fluctuations of tropomyosin. Sensing occurs by a weak binding mode in which the N terminus diffuses from actin monomer to monomer. Once specific cMyBP-C binding sites are encountered that allow subdomains C0 through C3 to bind more tightly, cMyBP-C can then effectively shift the tropomyosin equilibrium position from the blocked to the closed state, activating the thin filament. At high calcium (activating conditions), where calcium binding to troponin itself shifts tropomyosin from the blocked to the closed position, cMyBP-C can bind tightly to the thin filament and compete for myosin binding. It is important to note that our studies define the cMyBP-C N-terminal interactions with the thin filament but do not rule out potential interactions with the myosin head region as contributing to the thin filament sensitization. Regardless, our studies provide at least one molecular model for how cMyBP-C carries out its function within the confines of the sarcomere *in vivo*.

## Acknowledgements

We would like to thank Jeffrey Robbins and James Gulick at Cincinnati Children’s Hospital Medical Center for providing the expressed cMyBP-C N-terminal fragments and Greg Hoeprich, Lynn Chrin, and Christopher Berger at the University of Vermont for fluorescently-labeling and characterizing the N-terminal cMyBP-C fragments. These studies were supported by funds from the National Institutes of Health (HL126909, AR067279 to DMW; HL124041 to MJP) and the British Heart Foundation (FS/13/69/30504 to NMK and AVI). The authors declare no conflicts of interest.

AVI, SP, MP collected data. AVI, MP, DMW, NMK designed and conceived experiments. AVI, MP, DMW, NMK wrote the manuscript.

## Supplementary Information

To quantify the fluorescence intensity of a single fluorophore, we fitted a series of fluorescent spots seen at low calcium conditions where binding was sparse and therefore, more likely to derive from a single molecule. In addition, by keeping the frame rate and illumination power constant throughout all the acquisitions using Cy3-labelled cMyBP-C fragments, we ensured that the fluorescence emission from Cy3 remained approximately the same (59). These values for Cy3-C0C3 binding to thin filaments at low calcium (0.1 μM, pCa 7, Fig. 3a left panel) are shown as a histogram in Figure S1. This histogram was fitted to six Gaussian distributions using least squares in Microsoft Excel. We selected the maximum number of distributions for fitting by increasing the number of Gaussians until an additional Gaussian led to it being overlaid on top of another peak with no substantial improvement of fit. The intensity and the mean of the Gaussian distributions were fit independently while their standard deviation was constrained to one value for all the distributions, to reduce the number of degrees of freedom used in the fitting.

The mean values for each of the Gaussian components follow a linear relationship with the number of molecules within each fluorescence spot, with a slope of 372.4, corresponding to the intensity increase for the addition of another fluorophore (inset in Fig. S1). This value was used as a scaling term for the number of Cy3 molecules in all other conditions.

**Figure S1.**
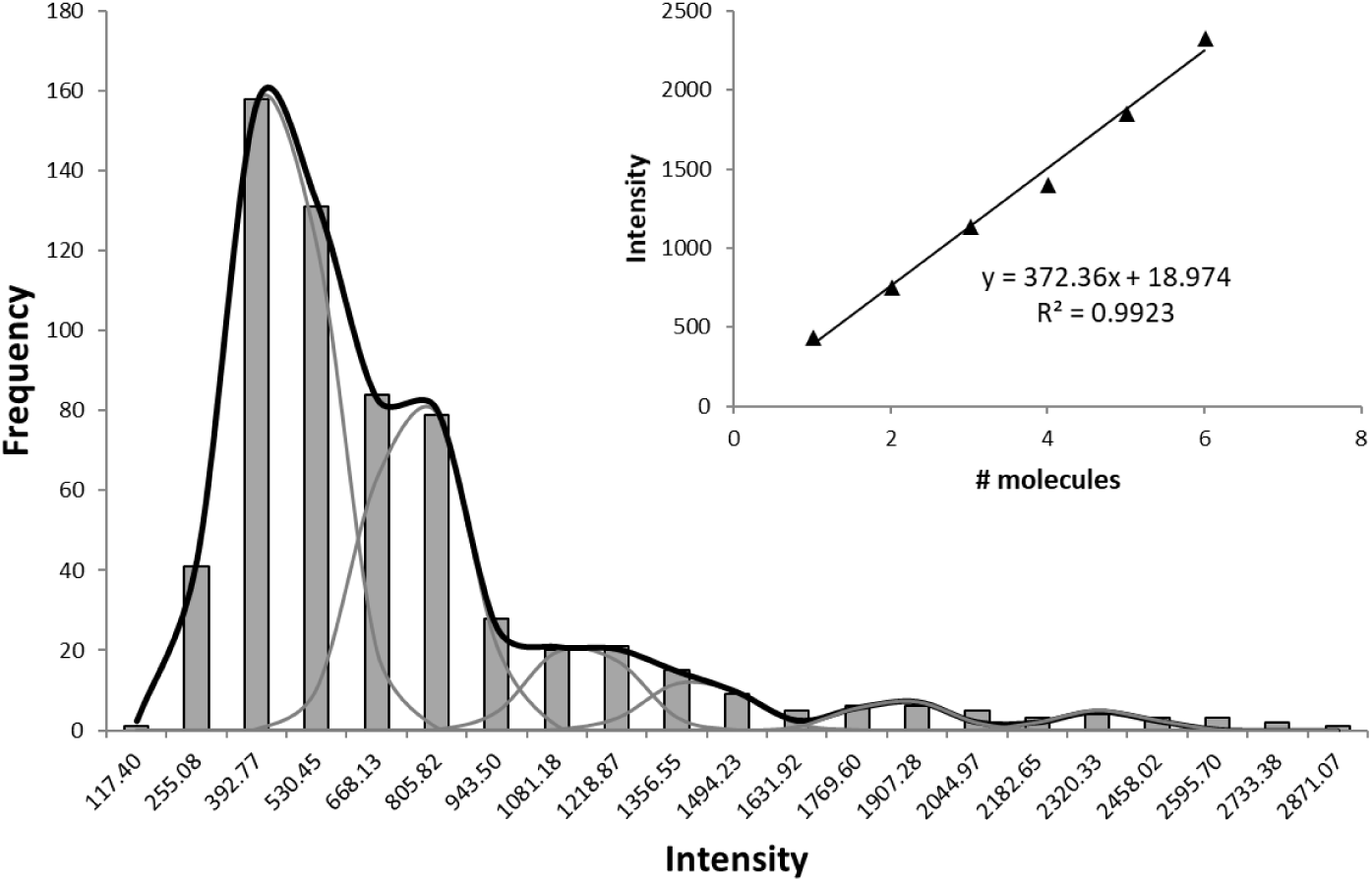
Determining the fluorescence intensity of a single Cy3-C0C3: a) A histogram of fluorescence spot intensities (bars) were fitted to a sum of Gaussian distributions (individual component Gaussians are shown in grey and the sum is the dark line) of fixed standard deviation. The inset shows the peak value for each Gaussian distribution plotted against a linear increasing scale, equivalent to the number of molecules present in that peak. The slope of this linear plot provides the fluorescence intensity of a single Cy3 molecule as 372.4 arbitrary units.

